# Generative Moment Matching Networks for Genotype Simulation

**DOI:** 10.1101/2022.04.14.488350

**Authors:** Maria Perera, Daniel Mas Montserrat, Míriam Barrabés, Margarita Geleta, Xavier Giró-i-Nieto, Alexander G. Ioannidis

## Abstract

The generation of synthetic genomic sequences using neural networks has potential to ameliorate privacy and data sharing concerns and to mitigate potential bias within datasets due to under-representation of some population groups. However, there is not a consensus on which architectures, training procedures, and evaluation metrics should be used when simulating single nucleotide polymorphism (SNP) sequences with neural networks. In this paper, we explore the use of Generative Moment Matching Networks (GMMNs) for SNP simulation, we present some architectural and procedural changes to properly train the networks, and we introduce an evaluation scheme to qualitatively and quantitatively assess the quality of the simulated sequences.

## I. INTRODUCTION

Genomic studies increasingly make use of large datasets and biobanks composed of thousands to hundreds of thousands of individual sequences. However, there are several challenges when dealing with such personal genomes: first, privacy restrictions can limit the scope of data sharing and access. Second, some of these datasets are very large introducing difficulty in storage, transfer, and processing. Finally, many of these datasets are highly imbalanced with extreme overrepresentation of some population groups (commonly European-descent individuals), and underrepresentation of others, leading to models that perform poorly when faced with individuals from underrepresented population groups [1]. Effective simulation tools, such as generative neural networks, can help to mitigate some of these challenges. By sharing trained generative networks, new synthetic sequences can be created to train machine learning models without the need to share the original sequences, (which can be prohibitive due to their size or privacy). Furthermore, through conditional synthesis (generation of sequences given a desired phenotype or ancestry), bias within datasets or biobanks can be partially removed by augmenting datasets with simulated samples from underrepresented groups.

Modern biobanks used in GWAS and similar studies typically include sequences of single nucleotide polymorphisms (SNPs). SNPs are the positions (less than 1%) within the human genome that are known to change between individuals. These positions (when biallelic) can be codified in a binary system, with 0’s denoting the common variant, and 1’s the minority variant. The nature of the sequences needs to be taken into account when designing generative networks: SNP sequences are sparse (they include many more 0s than 1s), high-dimensional, and commonly lack repetitive motifs. This completely differs in nature from other common signals, such as images or audio. The frequency distribution and correlations between SNPs vary across population groups, which can cause the techniques and treatments developed using data from one group not to generalize well to other population groups.

In this paper, we make use of a Generative Moment-Matching Network [2] to generate realistic genotype sequences given a desired ancestry. GMMNs, described in more detail in the following sections, are able to generate synthetic data that match the statistical moments of real data. This method has the same intent as other generative approaches, such as Generative Adversarial Network (GAN) [3] or Variational Autoencoders (VAEs) [4]. However, GMMNs can be easier to train than GANs or VAEs. Training a GAN requires solving a commonly difficult min-max optimization problem between two networks: a generator, and a discriminator. Similarly, a VAE tries to minimize reconstruction error and a KL-Divergence with two networks: an encoder and a decoder (generator). While such approaches require jointly training two networks, GMMNs require only training one unique network (the generator), which can lead to lower memory and computational requirements. In addition, GMMNs do not need to observe the data directly (as GANs or VAEs do), but instead can observe “sketches” (e.g. random feature descriptors [5]) that capture the statistical properties of the database as a whole. This allows for a privacy-friendly setting and permits a more accessible data sharing framework.

The contributions of this work are as follows: we introduce two modifications to the GMMN framework, a binarization step, and an iterative re-start of random projections, both described in section 3, that allow for a correct training and sequence simulation. Additionally, we propose two evaluation schemes: a qualitative evaluation through dimensionality reduction techniques, and a quantitative evaluation through supervised machine learning based classifiers.

## II. RELATED WORK

Generating realistic synthetic genotype sequences remains a challenge, as there are several evolutionary (selection and mutation) and demographic forces (drift) influencing genomes. Accurate simulation models should provide realistic allele frequency patterns and linkage disequilibrium (LD) profiles. There is an extensive literature of simulation methods based on evolutionary models such as the Wright-Fisher model [6]. Such techniques simulate recombination and create synthetic descendants by using available genetic sequences as ancestors. However, these models ignore evolutionary forces including selection and mutation. Coalescent simulation [7], [8] is a standard procedure to generate artificial genotypes under various models, usually seen as an extension of the classic Wright-Fisher model. Examples of such include Hudson’s ms program [9] that allows for mutation and migration within subpopulations, and ms reimplementations with improvements for modern datasets such as msprime [10], mbs [11], msms [12]. Because these models simplify reality, each of them has their own limitations, as the goal of any simulation tool is to find a compromise between the accuracy and efficiency [13].

Recently, new data-driven deep learning based simulation procedures have been introduced into the field – without defining an explicit evolutionary and demographic model. Connectionist approaches rely on learning models capable of generating realistic genotypes. The first technique to simulate SNP sequences from a population-based perspective was a class-conditional VAE-GAN by Montserrat et al. [14], which showed success on training local ancestry inference techniques. Yelmen et al. generate realistic surrogates of real genotype snippets with GANs and Restricted Boltzman Machines (RBM) [15]. Battey et al. tested VAE for genotype simulation and showed scaling issues of the method [16]. More recently, Geleta et al. introduced an ancestry-conditional VAE model for realistic genotype simulation [17], [18] with high-fidelity with respect to real genotype entropy variation and LD patterns. Generative models, additionally, provide a means for estimating meaningful low-dimensional representations which can be used for genotype imputation, classification, and visualization [17]. While being promising, such generative models do not provide extensive evaluation procedures to properly determine the quality of the simulated sequences. Some methods adapt a utility-preserving evaluation [14] where the quality of the sequences is assessed by their capacity to train neural networks in downstream task. Other methods use first order statistics or visualizations with a limited amount of dimensionality reduction techniques [15], [17]. Therefore, a proper procedure for simulation quality evaluation is still required.

## III. PROPOSED METHOD

### A. Generative Moment Matching Networks

GMMNs [2] are deep generative models able to generate new samples that statistically resemble the training samples. Such networks learn a mapping 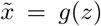 from a known distribution (typically Gaussian) into the data space; new samples are generated by sampling random vectors from 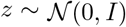 and processing them through the network. GMMNs are trained by minimizing Maximum Mean Discrepancy (MMD) criterion, which is a frequentist estimator that tries to determine if two sets of samples, 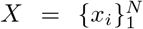 and 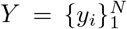, (in this case real and fake sequences) come from the same or different distributions. Specifically, MMD can be represented as the mean squared difference between two statistics:

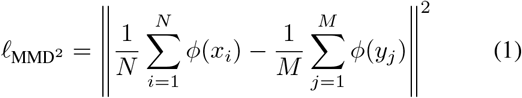

where 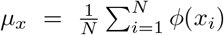 is a statistic, or “sketch”, computed on the samples *X*. If the statistics *μ_x_* and *μ_y_* are similar, then the two datasets are likely to come from the same distribution.

Taking *ϕ* as the identity function, the statistic *μ_x_* is equivalent to the sample mean, which for genotype sequences represents the frequency of each SNP. Other choices of *ϕ* can be used to capture higher order moments of the data distribution. Namely, as specified in [2], the kernel trick can be applied to obtain an infinite dimensional feature space that can capture all moments of the data. However, this requires computing a pairwise kernel distance with every training sample for each batch during the training process, which can be computationally expensive. Instead, we can use random features [5] to approximate any kernel with a finite number of dimensions. The random features are computed as *ϕ*(*x*) = *f*(*Wx*), where *f*(·) is a non-linearity, and *W* is a random projection matrix following some pre-specified distribution. When the number of features (dimension of *f*(*Wx*)) tends to infinity, the feature space tends to the reproducing kernel Hilbert space of the actual kernel. The type of kernel that is approximated depends on the distribution of *W* and the activation function applied. In this work we make use of ReLU random features which approximate the arc-cosine kernel [19].

Note that in order to train the network, the actual real genomic sequences do not need to be accessed, but just the statistic, or sketch, *μ_x_* is required. Typically, the sketch has a dimension *D* with *D* ≪ *NM*, where *N* is the number of samples, and *M* is the number of SNPs at each sequence. This provides a more compact representation of the data which is easier to share and to incorporate into privacy-preserving pipelines. However, this advantage is lost if the kernel trick is applied as in [2], where the actual samples are required to compute the pairwise kernel distances.

### B. Network Architecture

In this work, we perform ancestry conditional simulation by training a different GMMN for each ancestry group. Each GMMN is trained with a different sketch *μ* computed with the data of the specific ancestry group. This differs from other approaches were the ancestry label is provided as an extra input to the network [14]. While training a different network for each ancestry group can require a larger number of parameters (as many networks need to be trained), it provides a more flexible setting in scenarios where only the data of one ancestry group needs to be shared by sharing the specific trained network. For each ancestry-dependent network, we use the same architecture: a linear layer of dimension 5000 × 4096, followed by a ReLU activation and a batch norm, followed by another linear layer of dimension 4096 × 5000, finishing with a binary quantizer. Each network is trained with the MMD loss with their respective sketch vectors, where the sketch of the real data is pre-computed to save computing time.

**Fig. 1:**
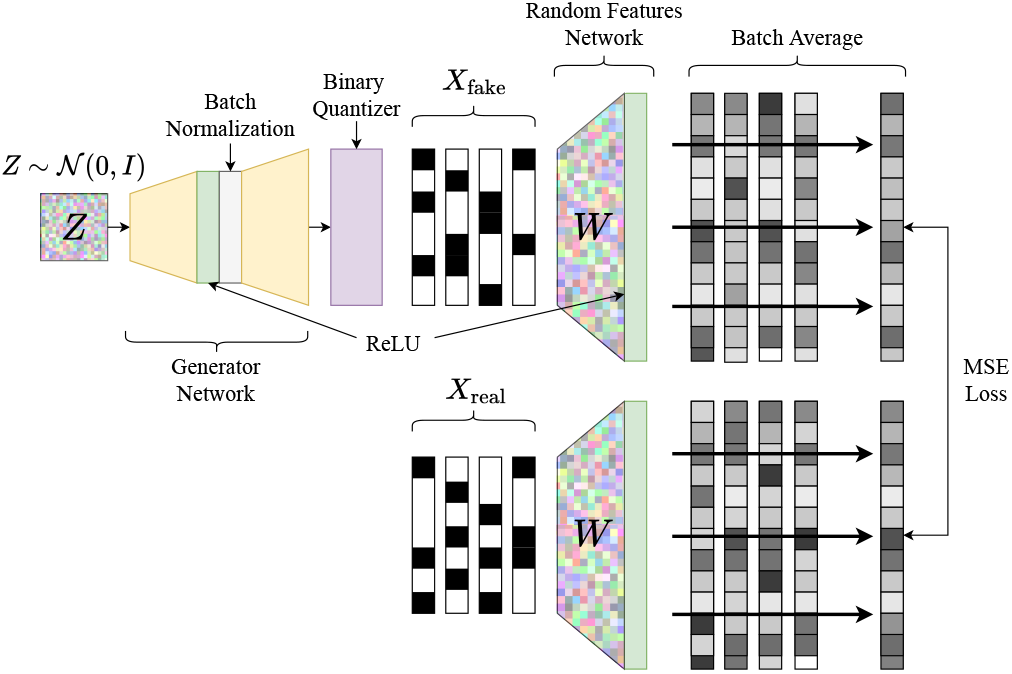
GMMN with Random Features.

The random features are implemented with a random linear layer of dimension 5000 × 50000, sampled from a Gaussian distribution, followed by a ReLU activation function. Note that the random linear layer that computed the random features is not trained.

### C. Output Binarization

Typically, generative networks output the simulated values through a linear or sigmoid layer. However, when simulating SNP sequences, only binary values are desired. Furthermore, binarizing the data through direct thresholding is a non-differentiable operator and does not allow the MSE loss signal to backpropagate through the network. Therefore, a differentiable alternative is required. In this work we make use of the same approach as in [20] where a hard threshold is applied in the forward pass (leading to binary simulated sequences), but the backward pass is approximated as if a scaled and shifted tanh operator was used during the forward pass.

### D. Random Features Re-start

In order to improve the quality of the simulation, we propose the use of an iterative re-start strategy of the random projections and sketch vectors. After a fixed number of weight update steps or number of training epochs, the random weights of the random feature projections are resampled and the sketch vector of the training dataset is recomputed. This allows us to improve the accuracy of the simulation and avoids having the network overfit on the specific features and statistical moments captured by the sketch vector. Note that a sketch with much larger dimensionality would allow for the same level of simulation accuracy, however, a larger dimensionality of the sketch requires a larger number of features to be computed at every batch leading to a slower training procedure. By performing re-sampling we obtain networks that capture rich features without the need of computing very high-dimensional features at every batch.

### E. Privacy And Data Sharing

Because access to the data is not required to train the network, and only access to the sketches *μ* are needed, our approach allows for a privacy-friendly setting with easy data sharing. By simply sharing the sketch vectors and the random projection matrices, different users can simulate new data by training GMMNs. Furthermore, as shown in [21] some computations can be already performed directly with the sketches, without the need of simulation of actual sequences. However, the sharing of sketches and/or trained networks, does not provide any strong guarantees on the privacy of the sequences used for training. Paradigms such as differential privacy can be incorporated easily into this framework in order to provide strong theoretical privacy guarantees, as it has been explored in [22], but we leave the exploration of its applications in genetic sequences as future work.

The use of different sketches for each ancestry group, and the re-start of random projection and sketches, can be seen as reflecting a real scenario where incoming available data comes from different sources (hospitals, academic labs, companies) each capturing populations from different countries and regions.

### F. Dimensionality Reduction for Qualitative Evaluation

When simulating natural signals such as images or audio, visual or auditory inspection can be used to evaluate the quality of the generated data. However, the high-dimensional nature of genomic data does not allow for an effective evaluation of simulated sequences through direct subjective inspection. Therefore, dimensionality reduction of the sequences is required in order to perform visual evaluation. Principal Components Analysis (PCA) has already been used in previous works such as [17] to visualize the synthesized sequences. We extend this approach into a more generic framework: we suggest that any dimensionality reduction technique that allows for projecting new unseen samples can be used to perform qualitative evaluation by first learning a projection into a lower dimensional space (usually 2 or 3 dimensions) with the real data, and later projecting the synthetic data using the learnt projection. If the projection of the synthetic data differs from the projection of the real data, that prima facie indicates that the simulation is not accurate. If the projections are not distinguishable between real and fake data, it is a sign that the simulation might be accurate. Visual inspection of dimensionality reduction projections is of course not enough to confirm the quality of the simulated data, and quantitative techniques are also required. Some dimensionality reduction techniques that can be used for such purpose include PCA, Linear Discriminat Analysis (LDA), Independent Component Analysis (ICA), Autoencoders (AE), ISOMAP [23], Parametric Uniform Manifold Approximation and Projection (UMAP) [24], and many others. In this work we showcase the use of PCA, ISOMAP, and Parametric UMAP. When applied to genetic data, most of the previously mentioned techniques will generate clusters according to the population structure of the data which is commonly correlated with the geography.

### G. Supervised Discriminators for Quantitative Evaluation

In this work, we explore the use of the accuracy of classifiers trained to distinguish between real and fake sequences as a quantitative measure of the quality of the simulation. Intuitively, if the simulation method is accurate, the classifiers should not be able to discriminate between real and fake samples and should obtain a classification close to 50% (random chance). In practice, many classifiers can memorize the training data obtaining a 100% classification accuracy within the training set. Therefore, we make use of the classification accuracy within a test set (sequences unseen during the training of the classifier) as an evaluation metric for the simulation method. Note that these classifiers have a similar role as the discriminator in GANs [25], but instead of providing a training gradient (as in GANs), they act as evaluation measures of the simulation quality. In fact, such classifiers can be used to evaluate the quality of any type of simulator, not necessarily based on Neural Networks.

We use Logistic Regression [26], K-Nearest Neighbors Classifier (KNN) [27] and Multi-Layer Perceptrons [28] for the evaluation task. We simulate the same numbers of synthetic sequences per ancestry as real sequences available. We randomly select 67% of both real and synthetic samples to comprise the training set and use the remainder as the testing set. We use five different random splits of the dataset and perform multiple times training and validation of the machine learning classifiers. Finally, we compute the average performance of each model and use the standard deviation to measure the variation in the accuracy results.

## IV. EXPERIMENTAL RESULTS

We use a subset of the dataset presented in [29]. This dataset includes human labeled genomic sequences from several publicly available databases. Our dataset contains 5000 SNPs from chromosome 22 codified in a binary format for 1683 individuals. Specifically, it includes a total of 358 Europeans (EUR), 403 Africans (AFR), 486 East Asians (EAS), and 436 South Asians (SAS) individuals.

We train multiple GMMNs to explore (1) the effect of replacing the sigmoid by a differential quantizer (binarization layer), (2) of using ReLU features instead of the mean of the dataset (*ϕ* = Identity), and (3) of re-starting random features during training. In table I, we present the classification accuracy (real vs fake) of the 3 discriminators: logistic regression, k-NN, and MLP. Note that simple classifiers (logistic regression and k-NN) can be fooled easily, leading to a discriminative accuracy close to random chance (50%), while the MLP obtains a better discriminative accuracy. Because some classifiers discriminate better than the others, we report the worst case scenario as the classification accuracy furthest from random chance (50% accuracy) within the Δ = |0.5 – *Acc*| column. A setting where at least one of the classifiers can classify real vs fake with 100% accuracy, will lead to Δ = 0.5, while a simulator that can fool all classifiers, will lead to Δ = 0.0. Therefore, a good simulation technique will have a Δ close to 0, while a bad one will have a Δ close to 0.5.

**TABLE I:**
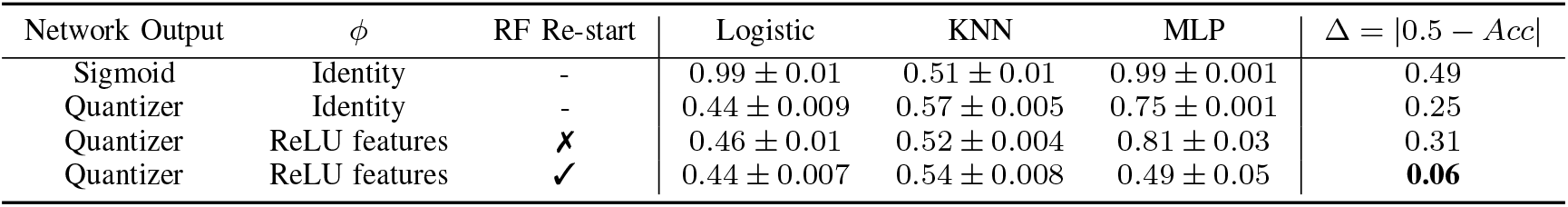
Mean and standard deviation of accuracies of the supervised discriminators with different GMMN configurations.

As shown in the first two rows of table I, including a differentiable binarization step instead of a sigmoid layer, leads to a significant improvement in the simulation quality (Δ decreases from 0.49 to 0.25). This can be observed in figure 2, where the PCA of simulated data is shown. Subfigure b) and c) shows the simulated data after and before thresholding of the sigmoid layer and d) shows the simulated data using the quanitzer layer. The real and fake data can be visually discriminated if a sigmoid layer is used. The use of ReLU features (instead of an identity mapping) might not lead to improvements on the simulation accuracy as shown in the second and third row of table I. Due to the stochastic nature of random features, some instances of the random mapping might not capture a correct set of features. However, by including random re-starts (last row of table I) the simulation accuracy with random features is highly increased obtaining sequences that practically fool all the trained discriminators. Finally, in figure 3 we show visualizations of UMAP and ISOMAP using our best configuration of GMMNs (Quantizer + Random Features + RF re-start), and we show that no apparent differences between real and simulated sequences are present.

**Fig. 2:**
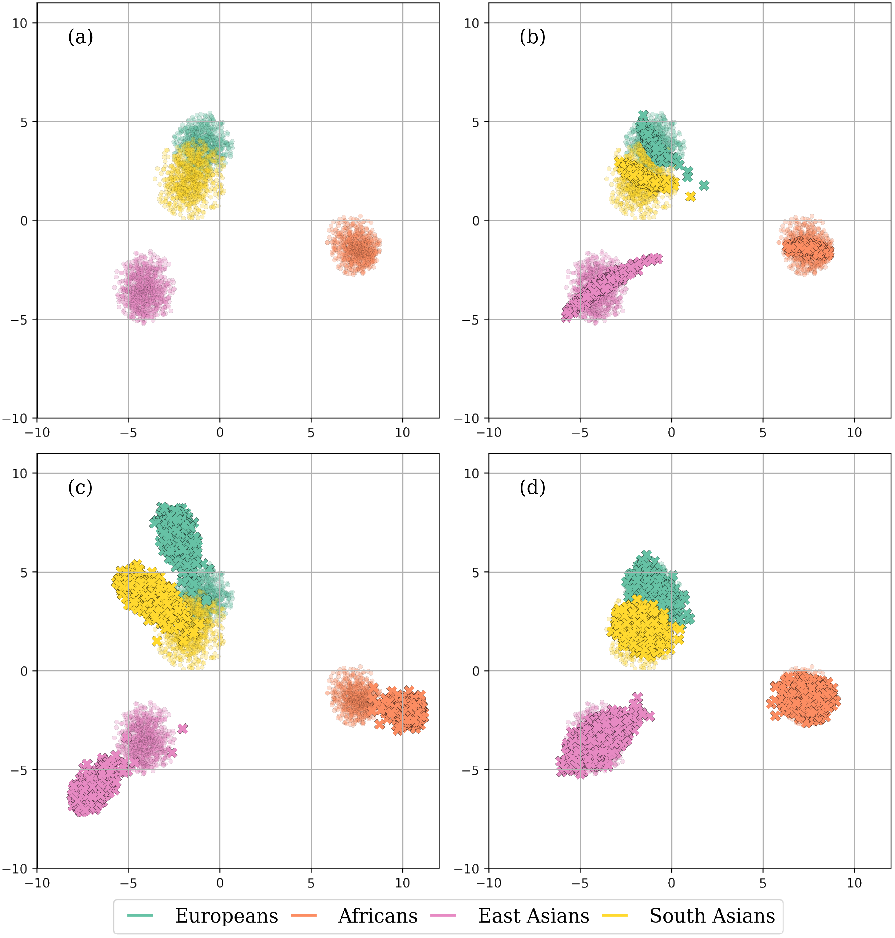
PCA of: (a) real data and generated data (b) with sigmoid output, (c) with sigmoid + threshold output, and (d) with binary quantizer output. Real data PCA is plotted in the background in b, c, and d.

**Fig. 3:**
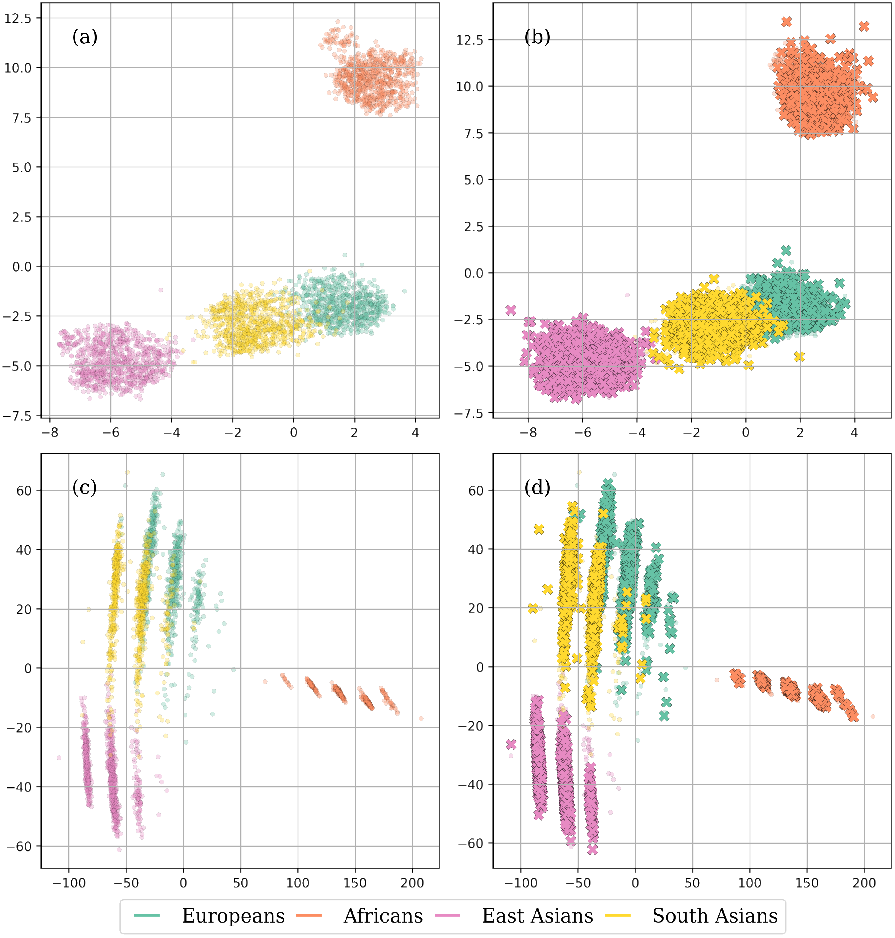
UMAP of (a) real data and (b) generated data. ISOMAP of (c) real data, and (d) generated data.

## V. CONCLUSIONS

In this work, we showed that GMMNs are able to successfully generate new, realistic samples given a desired ancestry and that simulation accuracy can be improved by using differentiable binarization and restarting the random features during training. Furthermore, we showed how evaluation of the simulated data can be performed with dimensionality reduction techniques and supervised classifiers. Finally, we discuss the potential of the presented framework to become a useful tool for privacy-preserving data sharing mechanism for modern day biobanks, and most importantly, a way to partially mitigate population bias within datasets.

